# Medroxyprogesterone reverses tolerable dose metformin-induced inhibition of invasion via matrix metallopeptidase-9 and transforming growth factor-β1 in KLE endometrial cancer cells

**DOI:** 10.1101/596056

**Authors:** Dong Hoon Suh, Sunray Lee, Hyun-Sook Park, Noh Hyun Park

## Abstract

This study was performed to evaluate the anticancer effects of tolerable doses of metformin with or without medroxyprogesterone (MPA) in endometrial cancer cells. Cell viability, cell invasion, and levels of matrix metallopeptidase (MMP) and transforming growth factor (TGF)-β1 were analyzed using three human endometrial adenocarcinoma cell lines (Ishikawa, KLE, and USPC) after treatment with different dose combinations of MPA (0, 10 μM) and metformin (0, 100, 1000 μM). Combining metformin (0, 100, 1000 μM) and 10 μM MPA induced significantly decreased cell viability in a time- and dose-dependent manner in Ishikawa cells, but not in KLE and USPC cells. There was no dose- or time-dependent cell growth inhibition, or positive western blot results for the expression of progesterone receptors and phospho-AMPKa, following treatment with any combination of metformin (0, 100, 1000 μM) and 10 μM MPA in KLE and USPC cells. In KLE cells, metformin treatment alone significantly inhibited cell invasion in a dose-dependent manner (1.31±0.05, 0.94±0.04, 0.83±0.05 at 0, 100 μM, 1000 μM, respectively; p<0.0005). The inhibitory effect of metformin was reversed to create a stimulating effect when metformin was combined with 10 μM MPA (1.10±0.05, 1.42±0.18, 1.41±0.26 at 0, 100, 1000 μM, respectively; p<0.005). MMP-9 and TGF-β1 showed similar trends in terms of cell invasion in KLE cells. In conclusion, the anti-invasive effect of metformin in KLE cells was completely reversed to the state of no treatment by the addition of MPA; this might be mediated through MMP-9 and TGF-β1. Our study suggests the possibility of these combinations doing harm, rather than good, under some conditions.

## Introduction

Uterine corpus cancer was found to be the 4th most common cancer in women in 2018; the estimated number of new cases was 63,230, accounting for 7% of all new cancer diagnoses in women (1). Endometrial cancer constitutes the majority of uterine cancers, excluding uncommon subtypes of stromal or mesenchymal sarcomas, which account for approximately 3% of all uterine cancers. Despite multimodal treatment approaches, type I poorly differentiated endometrioid adenocarcinoma and type II cancers, including uterine serous papillary cancer (USPC) without estrogen receptor (ER) and progesterone receptor (PR) expression, have very poor prognosis unlike type I well-differentiated endometrioid adenocarcinoma, which expresses ER and PR. Among the systemic hormonal therapies considered for recurrent, metastatic, or high-risk disease, progestin is the most commonly used, mainly in the form of medroxyprogesterone acetate (MPA). However, clinical guidelines recommend that MPA may only be used for lower-grade endometrioid histology. This is based on previous reports that the highest response rates were noted in low-grade, ER-positive tumors of up to 55% (2, 3). In addition, long-term continuous use of progestin was known to cause a loss of effect of PR activation (2). Therefore, development of a new treatment strategy for groups of cancer with poorer prognosis is urgent.

Recently, metformin, an oral biguanide anti-diabetic drug for type 2 diabetes, was shown to have significant anticancer activity and considered a novel treatment option through drug repositioning (4), including for endometrial cancer (5-7). However, it should be noted that almost all previous studies were conducted with supra-pharmacological concentrations (doses) of metformin, that is, 10–100 times higher than maximally achievable therapeutic concentrations found in patients with type 2 diabetes mellitus (8). Such levels exceed the maximum dose that could cause lactic acidosis, one of the most serious side effects of metformin. Any anticancer effect of metformin should be studied only in the condition of achievable therapeutic concentrations (8, 9).

Another approach for the development of novel anticancer drug regimens is the use of drug combinations. Although hormonal therapy is currently recommended only for lower-grade endometrioid histology in clinical guidelines, there is evidence suggesting several anticancer mechanisms of progestational agents, which could show significant effects in poorly-differentiated endometrioid adenocarcinoma, as well as USPC (2, 10, 11), particularly when combined with metformin (12).

The purpose of this study was to evaluate the anticancer effect of tolerable doses of metformin alone or with MPA in endometrial cancer cells.

## Materials and Methods

### Cell Cultures

Three human endometrial adenocarcinoma cell lines were used: Ishikawa (type I well-differentiated, ER+/PR+), KLE (type I poorly differentiated, ER-/PR-), and USPC (type II serous papillary carcinoma, ER-/PR-) (13). Ishikawa cells were purchased from the Japanese Collection of Research Bioresources cell bank and maintained in Dulbecco’s Modified Eagle Medium (DMEM) (Life Technologies) containing 10% fetal bovine serum (FBS) (Hyclone, Logan), 50 μg/ml streptomycin, and 50 U/ml penicillin. KLE and USPC cells were obtained from the American Type Culture Collection (ATCC, USA). KLE was cultured in DMEM/F12 medium (Life Technologies, CA, USA) with 10% FBS and 0.5% P/S and USPC cells were cultured in RPMI1640 medium (Life Technologies, CA, USA) with 10% FBS, 2 mM/L glutamine, and 0.5% P/S. All the cells were cultured in an incubator at 37°C under a humidified atmosphere containing 5% CO2.

### Dose setup of metformin treatments

Tolerable doses of metformin (500 mg twice/day) could achieve a plasma concentration of around 1 mg/L (14). Although the maximal approved total daily dose of metformin for treatment of diabetes mellitus is 2.5 g (35 mg/kg body weight)(8), slow but progressive increase of fasting lactic acid levels during metformin treatment with multiple doses from 100 to 850 mg twice a day suggested that; the higher dose of metformin, the higher risk of lactic acidosis (14). Therapeutic plasma concentrations or ranges of metformin measured in previous studies of type 2 diabetes ranged from 0.129 to 90 mg/L (9). Therefore, 1 mM (129.2 mg/L) was set as a maximal concentration of metformin for our experiment, enabling the maximum achievable plasma concentration in a clinical setting without the risk of lactic acidosis.

### Cell counting and cell survival analysis

To increase cell growth rate, all cells were seeded in 12-well plates (Corning Life Sciences, NY, USA) at 10,000 cells/cm^2^, and the cell number was counted at 24-hour intervals until 96 hours. For cell counting, the medium was removed from the cell culture plates, washed twice with phosphate buffer saline (PBS), and then treated with 0.25% trypsin for 5 min at 37 °C. The trypsin-treated cells were collected in a 15 ml tube, washed twice with the culture medium, and counted three times using the Adam-MC automatic cell counter (NanoEntek, Korea). Viable cells were more accurately measured using an advanced image analysis program of Adam-MC cell counter.

Cell survival analysis was performed to investigate the effects of metformin (Sigma-Aldrich) and/or MPA (Sigma-Aldrich) on endometrial cancer cell lines. Cells were seeded into a 96 well plate (Ishikawa 5*10^4^/cm^2^, KLE 2*10^4^/cm^2^ and USPC 3*10^4^/cm^2^). The next day, cells were treated with 100 μM and 1 mM of metformin, with or without 10 μM of MPA. Then, survival rates of cells were analyzed after 24 hours and 48 hours of drug treatment using a 3- [4,5-dimethylthiazol-2-yl]-2,5 diphenyl tetrazolium bromide (MTT) assay kit (DoGen Co., Korea). The assay was performed according to the supplier protocol (http://www.dogenbio.com/shop/item.php?it_id=1490923054).

### Western blot

The proteins collected from each cell sample were quantitated, subjected to 12% SDS-PAGE, and then transferred to a nitrocellulose membrane. The membrane was subjected to blocking in PBS, containing 0.1% Tween20 (Sigma) and 5% skim milk (Invitrogen), probed with primary antibodies, progesterone receptor-B (C1A2), AMPKα (Thr172), phospho-AMPKα (Thr172) (Cell Signaling, MA, USA), and ErbB2 (Abcam, Cambridge, UK), and then reacted with peroxidase conjugated secondary antibody (Jacson Immuno Research, USA). Finally, target bands were visualized using SuperSignal chemiluminescent (ThermoFisher Scientific, MA, USA).

### Cell invasion assay and ELISA

To perform invasion assays, we first coated matrigel (BD science, CA, USA) on a transwell membrane with 8 μm pores (Corning Life Sciences, NY, USA) at 37°C for 2hours, and seeded 8*10^4^ cells/cm^2^ into the transwell membrane. The next morning, the cells were starved for 2 hours in culture medium without FBS and the outside of the transwell was replaced with a 5% charcoal strip FBS (Life Technologies) containing medium to induce invasion for 24 hours in a 37°C, humidified atmosphere containing 5% CO2; anticancer drugs were treated according to the conditions at the time of the exchange of the medium. The next day, all the cells in the transwells were removed, and the transwells were inverted to stain the transferred cells with 0.2% crystal violet. The stained cells were de-stained with 2% SDS and absorbance was measured at 560 nm.

For quantitative analysis of cell migration related proteins, the secretion levels of Matrix metallopeptidase (MMP)-2 and -9 (R&D system, MN, USA) and transforming growth factor (TGF)-β1 (R&D system, MN, USA) were checked using ELISA kits. First, all cells were plated at 9*10^4^ cells/cm^2^ into a 24 well plate (Corning Life Sciences, NY, USA) and starved for 2 hours in culture medium without FBS. The anticancer drugs were treated according to the conditions while the culture medium was exchanged with the complete medium. After 24 hours, the cultures were collected without cells and analyzed. An ELISA was performed according to the supplier protocol (https://www.rndsystems.com/).

### Statistical analysis

All statistical analyses were performed using GraphPad PRISM (GraphPad Software Inc., CA, USA). The differences between the two groups were determined using a two-tailed unpaired Student’s t-test. A p value <0.05 indicated statistical significance.

## Results

### Cell growth and growth inhibition by tolerable doses of metformin and MPA in endometrial cancer cell lines

We found that USPC cells had the fastest growth rate among the three endometrial cancer cells during 96-hour incubation, followed by Ishikawa and KLE cells (Fig 1A and 1B). The MTT assay showed that treatment with metformin alone at ≤ 1000 μM for 48 hours exerted significant inhibitory effects on the cell viability of Ishikawa, KLE, and USPC cells in a dose-dependent, but not in a time-dependent manner (Fig 1C-1E). In Ishikawa cells, a combination of metformin (0, 100, 1000 μM) and 10 μM MPA induced a significant decrease in cell viability in a time- and dose-dependent manner (Fig 1C). Addition of 10 μM MPA to metformin significantly inhibited cell viability compared to that by metformin alone at each dose (0, 100, 1000 μM), respectively, in Ishikawa, but not in KLE and USPC cells.

**Fig 1.**
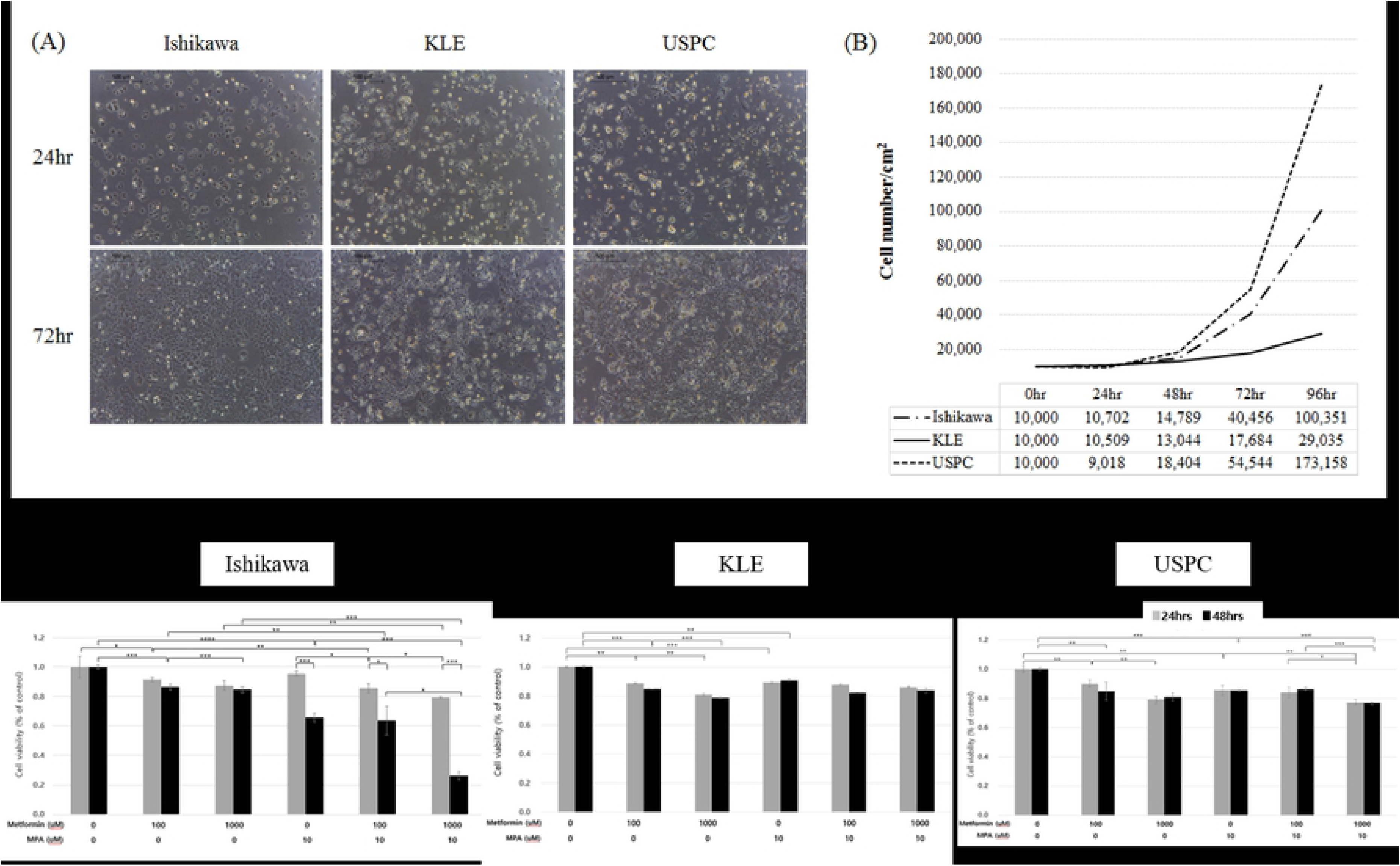
Cell growth and growth inhibition by metformin and medroxyprogesterone (MPA) in three endometrial cancer cell lines: Ishikawa, KLE, and USPC. (A) Cell morphology and number at 24 hours and 72 hours, (B) cell growth rate (0, 24, 48, 72, and 96 hour), cell viability after treatment of different dose combinations of MPA (0, 10 μM) and metformin (0, 100, 1000 μM) in Ishikawa (C), KLE (D), and USPC cells (E). *p< 0.05, **p<0.005, ***p<0.0005, ****p<0.00005

### Changes in expression levels of PR and AMPK by a tolerable dose of metformin and MPA in endometrial cancer cell lines

A significant level of endogenous expression of PR-B was found in Ishikawa cells but not in KLE and USPC cells (Fig 2). In Ishikawa cells, metformin treatment alone induced the expression of PR-B in a dose-dependent manner, whereas metformin combined with 10 μM MPA inhibited PR-B expression in a dose-dependent manner. Expression of AMPKα and its activated form, phospho-AMPKα (p-AMPKα), were inhibited by metformin treatment alone in a dose-dependent manner (0, 100, 1000 μM) in Ishikawa cells. However, p-AMPKα lost its dose-dependent pattern when Ishikawa cells were treated with a combination of metformin (0, 100, 1000 μM) and 10 μM MPA. In KLE and USPC cells, there were no significant changes in expression patterns of ErbB2, AMPKα, and p-AMPKα when treated with any of the doses of metformin and MPA. In USPC cells, the expression of p-AMPKα was stronger when both metformin and MPA were used than when metformin was used alone. However, there was neither dose dependency in expression patterns nor consistency with anti-proliferative effects (Fig 1E).

**Fig 2.**
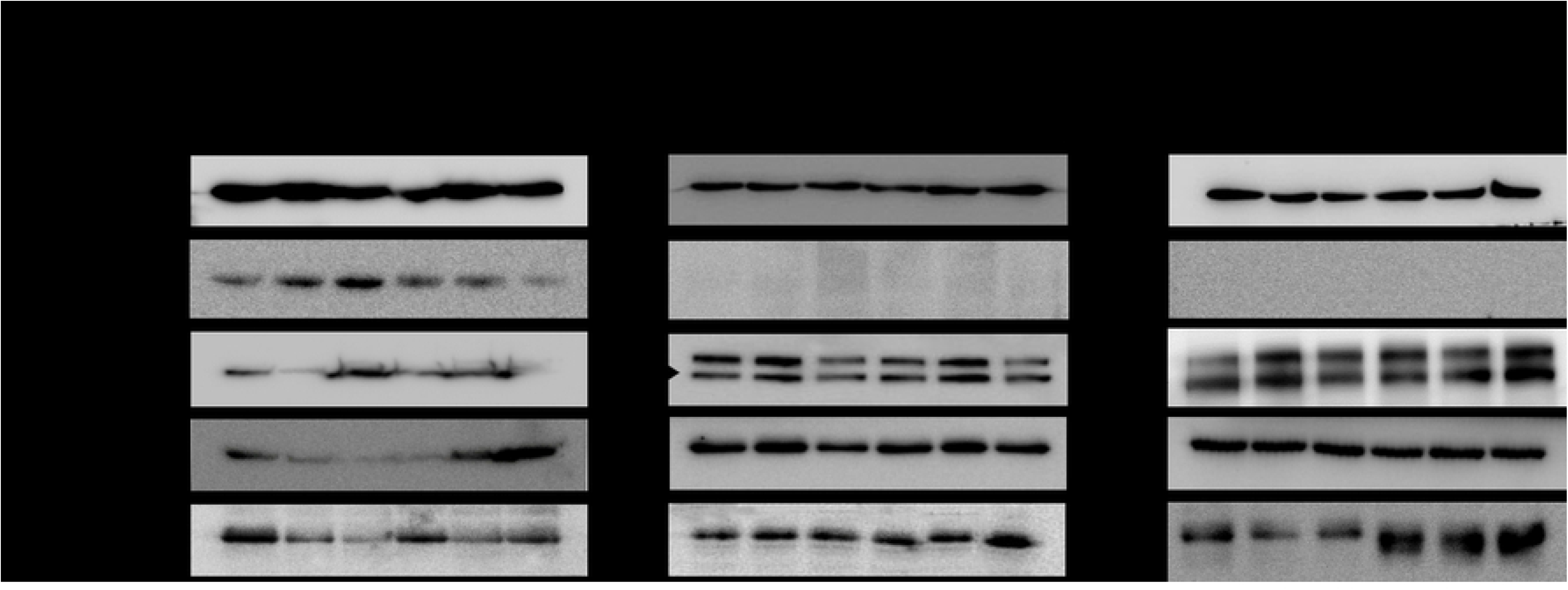
Expression of progesterone receptor B, ErbB2, AMPKα, and phospho-AMPKα in three endometrial cancer cell lines (Ishikawa, KLE, and USPC) according to treatment with different dose combinations of medroxyprogesterone (MPA) (0, 10 μM) and metformin (0, 100, 1000 μM).

### Inhibition and disinhibition of cell invasion by a tolerable dose of metformin with or without MPA in endometrial cancer cell lines

There was no dose- or time-dependent cell growth inhibition or positive western blot results for PR and p-AMPKα expression when any combination of metformin (0, 100, 1000 μM) and 10 μM MPA were used in KLE and USPC cells. We further performed an invasion assay (Fig 3A), which showed that metformin treatment alone did not induce any significant changes in cell invasion in Ishikawa and USPC cells (Fig 3B and 3D). In KLE cells (Fig 3C), however, metformin treatment alone significantly inhibited cell invasion in a dose-dependent manner (1.31±0.05, 0.94±0.04, 0.83±0.05 at 0, 100 μM, 1 mM, respectively; p < 0.0005). Treatment with MPA 10 μM alone significantly decreased the invasion of KLE cells compared to that of control cells (1.31±0.05 vs. 1.10±0.05; p<0.005). Interestingly, the inhibitory effect of metformin alone on cell invasion was reversed to have a stimulating effect when metformin was combined with 10 μM MPA (1.10±0.05, 1.42±0.18, 1.41±0.26 at 0, 100, 1000 μM, respectively; p < 0.005) (Fig 3C). In Ishikawa cells, by contrast, a combination of 10 μM MPA with metformin exerted a synergistic effect on the inhibition of cell invasion (0.93±0.05, 0.76±0.01, 0.69±0.01, at 0, 10 μM MPA alone, 100 μM metformin and 10 μM MPA; p<0.0005), although the synergistic effect disappeared at a metformin dose of 1000 μM (0.84±0.08) (Fig 3B). There was no significant effect of metformin and MPA combination on the invasion of USPC cells (Fig 3D).

**Fig 3.**
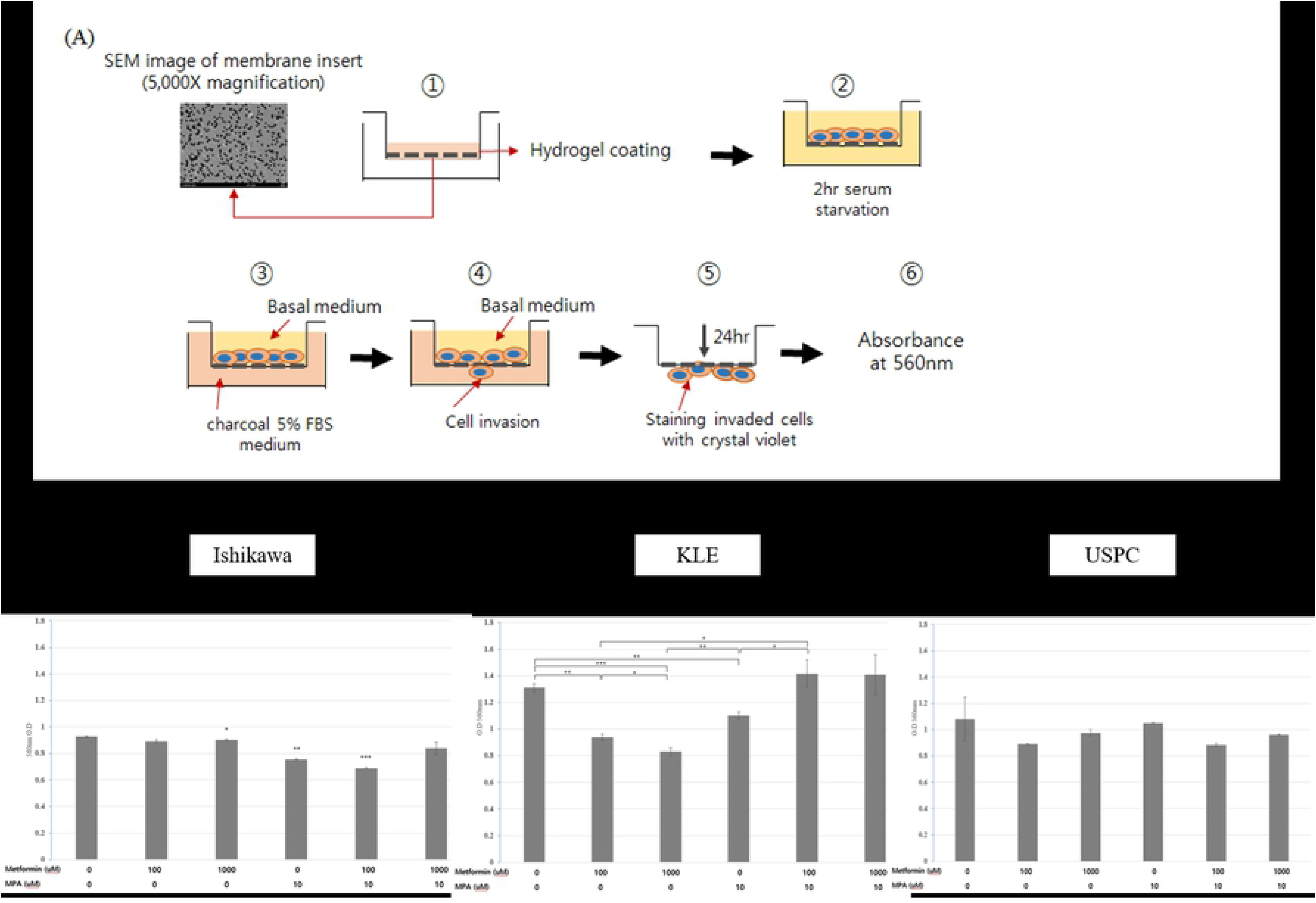
Invasion assay in three endometrial cancer cell lines (Ishikawa, KLE, and USPC). (A) The process of invasion assay, cell invasion after treatment of different dose combinations of medroxyprogesterone (MPA) (0, 10 μM) and metformin (0, 100, 1000 μM) in Ishikawa (B), KLE (C), and USPC cells (D). *p< 0.05, **p<0.005, ***p<0.0005

### MPA reverses tolerable dose metformin-induced inhibition of invasion via MMP-9 and TGF-β1 in KLE endometrial cancer cells

MMP-2 showed no significant changes in response to the treatments in all three cell lines (Fig 4A-4C). However, although decreased MMP-9 induced by metformin treatment alone was not statistically significant, MMP-9 showed the same trends of rise with cell invasion in KLE cells when treated in combination with metformin (0, 100, 1000 μM) and 10 μM MPA (3.99±0.90 for control, 5.83±1.04, 7.68±1.38, 8.05±2.09 ng/ml, respectively; p<0.05) (Fig 4E). Otherwise, there were no significant changes in MMP-9 expression in Ishikawa and USPC cells (Fig 4D and 4F).

**Fig 4.**
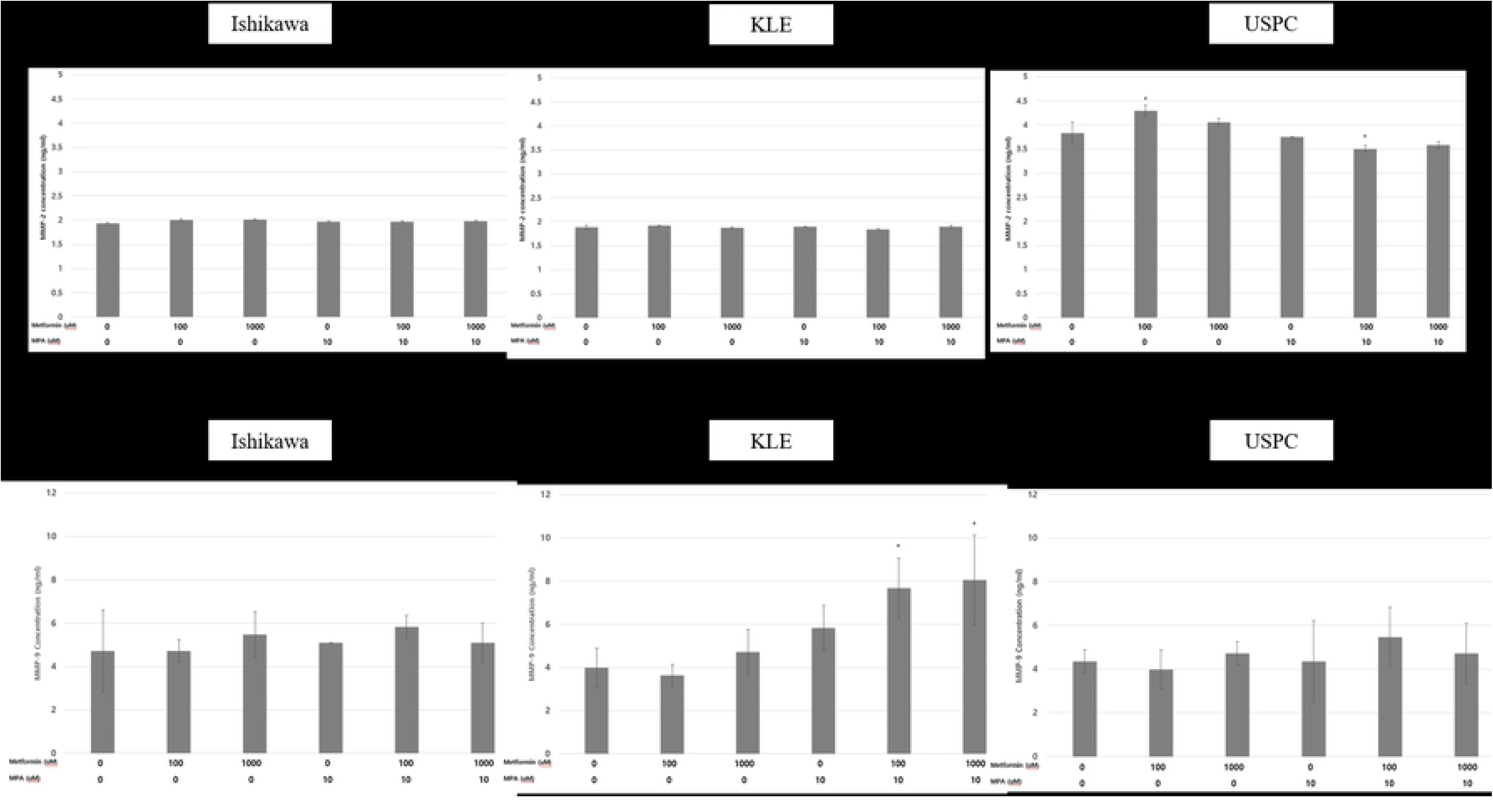
Matrix metallopeptidase (MMP)-2 (A, Ishikawa; B, KLE; C, USPC) and MMP-9 (D, Ishikawa; E, KLE, F, USPC) in three endometrial cancer cell lines after treatment with different dose combinations of medroxyprogesterone (MPA) (0, 10 μM) and metformin (0, 100, 1000 μM) *p< 0.05.

TGF-β1 also showed similar trends to MMP-9, which was in concordance with the change in cell invasion (Fig 5B). TGF-β1 secretion was significantly decreased when KLE cells were treated with 1000 μM metformin alone compared to that in control cells (62.76±2.18 vs. 54.19±3.60 pg/mL; p=0.024). Furthermore, TGF-β1 also exhibited the reverse pattern when treated with a combination of 1000 μM metformin and 10 μM MPA (62.76±2.18 vs. 77.52±5.95; p=0.016). There were no significant changes in TGF-β1 levels according to the treatments in Ishikawa and USPC cells (Fig 5A and 5C).

**Fig 5.**
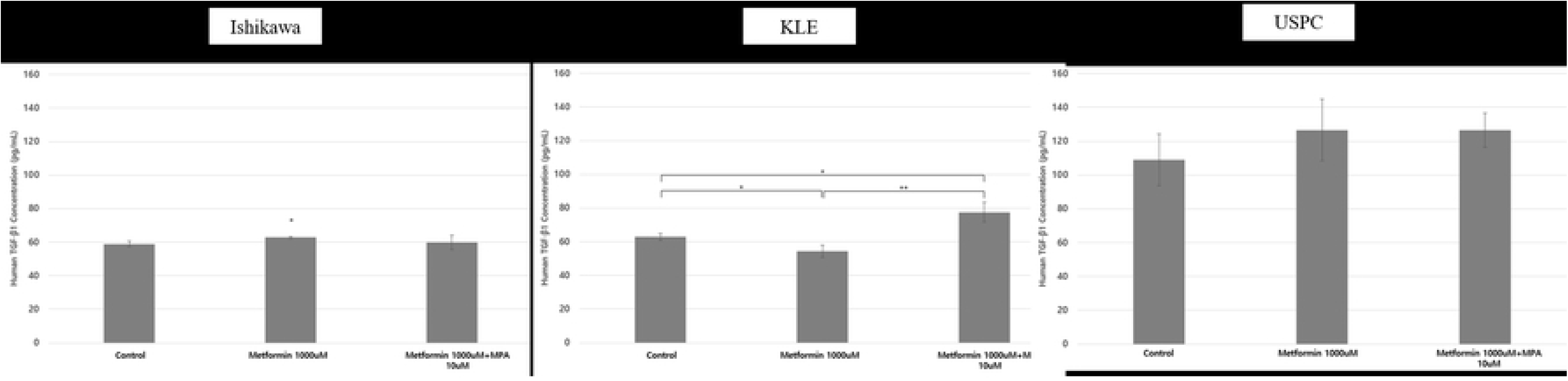
Transforming growth factor (TGF)-β1 in three endometrial cancer cell lines after treatment with different dose combinations of medroxyprogesterone (MPA) (0, 10 μM) and metformin (0, 1000 μM) in Ishikawa (A), KLE (B), and USPC cells (C). *p< 0.05, **p<0.005.

## Discussion

The principal finding of our study was that tolerable doses of metformin alone ≤ 1000 μM have anti-invasive effects on KLE cells, and the anti-invasive effect of metformin is even reversed by the addition of 10 μM MPA. Changes in the expression of MMP-9 and TGF-β1 were plausible mechanisms underlying these findings. We also showed that tolerable doses of metformin alone, ≤ 1000, μM inhibited cell proliferation of Ishikawa, KLE, and USPC cells in a dose-dependent manner. However, there was no additive or synergistic anti-proliferative effect of metformin ≤ 1000 μM and MPA co-treatment in KLE and USPC cells.

MPA is recommended as a fertility-preserving treatment for young endometrial cancer patients, as well as palliative treatment for terminally ill patients with hormone receptor-positive cancer, especially with PR; most of the MPA anticancer effects are known to act through the interaction with PR (15). However, response rates of MPA are unsatisfying because of the appearance of progesterone resistance; efforts were made to find an effective way to overcome this (12, 15, 16). Metformin was suggested to combine with MPA, based on several mechanisms of reversing progesterone resistance, mainly as a potent inhibitor of the PI3K-AKT-mTOR pathway by activating AMPK. However, there were two issues to be solved. One is the supra-therapeutic concentration of metformin. The activation of AMPK was almost always demonstrated at an unrealistically high supra-therapeutic concentration of metformin, considering the maximal dose in humans (without the risk of serious side effects) (16, 17). The other is that most of the findings were true only in Ishikawa cells, but not in other types of cells, for example, KLE (12). Therefore, we tried to find any anti-cancer effects of tolerable doses of metformin with or without MPA in endometrial cancer cells with non-favorable clinical behavior.

We found that metformin alone at ≤ 1000 μM significantly inhibited the proliferation of all three cell lines (Fig 1C-1E and 2). The anti-proliferative effect of metformin alone at ≤ 1000 μM in PR-positive Ishikawa cells might be mediated through PR-B. This is because the expression of PR-B but not of p-AMPK-α increased in a dose-dependent manner with metformin treatment (Fig 2). On the other hand, growth inhibition by low-dose metformin alone of KLE and USPC cells and the synergistic anti-proliferative effect of metformin in combination with MPA in Ishikawa cells were neither associated with PR-B nor with the AMPK-dependent pathway because there were no corresponding changes in their expression levels (Fig 2). The plausible mechanism underlying this synergistic anti-proliferative effect of the metformin and MPA combination on Ishikawa cells could include AMPK-independent pathways, including factors of the Rag family of GTPases, hypoxia inducible factor (HIF) target gene, and regulated in development and the DNA damage response I (REDD1) (17). Other studies confirmed that there was no significant increase in p-AMPKα expression at low doses of metformin, ≤ 1000 μM, in Ishikawa, KLE, and USPC cells (18, 19). In these studies, high dose metformin ≥10 mM was shown to be necessary to bring about a significant increase in p-AMPKα expression. As there were no significant changes in PR and p-AMPKα levels in KLE and USPC cells when metformin was combined with MPA (Fig 2), we moved our focus towards invasion; the plausible invasion mechanism was not related to the activation of AMPK.

It was interesting that the dose-dependent inhibitory effect of metformin ≤ 1000 μM on cell invasion was found only in KLE cells, but not in Ishikawa and USPC cells. It was even more interesting that the addition of MPA to metformin resulted in the opposite effects on cell invasion in the two different types of cells, i.e. stimulating (reversing metformin effect) in KLE and inhibitory (possible synergistic effect) in Ishikawa cells (Fig 3B and 3C). The finding of low dose metformin alone not conferring any change in invasion capability of Ishikawa cells was consistent with that in a study of de Barros Machado et al. (20). Even though we could not find a plausible molecular mechanism for the synergistic anti-invasive effect of the narrow dose window of metformin (0-100 μM) and MPA combination in Ishikawa cells (Fig 3B), it is notable that MMP-9 and TGF-β1 showed the same pattern of change to that of cell invasion only in KLE cells (Fig 4 and 5). MMP is known as one of the most essential proteases in the proteolysis of extracellular matrix proteins and plays important roles in invasion and migration (21). Unlike other relevant studies of MMPs in which the results were the same in MMP-2 and MMP-9 (21-23), the results of our study were specific to MMP-9, but not MMP-2. TGF-β has also been extensively studied as a crucial cytokine which promotes endometrial cancer invasion and metastasis via epithelial mesenchymal transition (21, 23, 24). MMPs and TGF-β form an interplay loop in which one might activate the other and vice versa, to facilitate tumor progression. The results of our study suggest that this loop could be inhibited by metformin at doses as low as less than 1000 μM, but completely disinhibited or even stimulated by cotreatment of MPA and metformin in KLE cells. Samarnthai et al. (25) reported the dualistic model of endometrial carcinoma, type I and type II, in terms of genetic changes and clinical behavior. KLE cells could be clinically characterized as type II cancer cells because of the aggressive behavior and poor outcomes, but also as type I due to the frequent PTEN and KRAS mutations and rare p53 mutation, which are typical in type I cancer.

Despite a small number of studies, so far, addressing the anti-invasive and/or anti-migratory effects of metformin in endometrial cancer cells, this study is, to the best of our knowledge, the first study which showed that the significant anti-invasive effect of a tolerable dose of metformin in KLE cells was completely reversed to the state of no treatment by the addition of MPA; these findings might be mediated through MMP-9 and TGF-β1. However, our study has some limitations in that metformin was not shown as an AMPK activator. However, there are a few studies which did not support metformin as a potent AMPK activator, not only in proliferation (18), but also in invasion (22). Some values were out of the expected ranges, for example, MMP-9 concentration after treatment with 1000 μM metformin only, which was expected to be lower than that of 100 μM metformin. Lastly, treatment with inhibitors such as GM6001 (an MMP inhibitor) could have made our results more confirmative.

Most studies on metformin and MPA in endometrial cancer have concluded that combining the two could be a potential therapeutic strategy for overcoming progesterone resistance (12, 15). However, our study suggests the possibility of the combination being harmful instead of beneficial in some conditions, especially in clinically highly aggressive but genetically type I cancer. Further animal studies are required to clinically confirm our study findings.

## Disclosure of conflict of interest

No potential conflict of interest relevant to this article was reported.

